# Identification of potent and safe antiviral therapeutic candidates against SARS-CoV-2

**DOI:** 10.1101/2020.07.06.188953

**Authors:** Xia Xiao, Conghui Wang, De Chang, Ying Wang, Xiaojing Dong, Tao Jiao, Zhendong Zhao, Lili Ren, Charles S Dela Cruz, Lokesh Sharma, Xiaobo Lei, Jianwei Wang

## Abstract

COVID-19 pandemic has infected millions of people with mortality exceeding 300,000. There is an urgent need to find therapeutic agents that can help clear the virus to prevent the severe disease and death. Identifying effective and safer drugs can provide with more options to treat the COVID-19 infections either alone or in combination. Here we performed a high throughput screen of approximately 1700 US FDA approved compounds to identify novel therapeutic agents that can effectively inhibit replication of coronaviruses including SARS-CoV-2. Our two-step screen first used a human coronavirus strain OC43 to identify compounds with anti-coronaviral activities. The effective compounds were then screened for their effectiveness in inhibiting SARS-CoV-2. These screens have identified 24 anti-SARS-CoV-2 drugs including previously reported compounds such as hydroxychloroquine, amlodipine, arbidol hydrochloride, tilorone 2HCl, dronedarone hydrochloride, and merfloquine hydrochloride. Five of the newly identified drugs had a safety index (cytotoxic/effective concentration) of >600, indicating wide therapeutic window compared to hydroxychloroquine which had safety index of 22 in similar experiments. Mechanistically, five of the effective compounds were found to block SARS-CoV-2 S protein-mediated cell fusion. These FDA approved compounds can provide much needed therapeutic options that we urgently need in the midst of the pandemic.

## Introduction

Novel coronavirus mediated disease (COVID-19) emerged as a major pandemic and has spread across the world in such a short span of time since Dec 2019. As of May 15, 2020, more than 4.5 million confirmed infections have been reported with approximately 308,000 deaths (WHO, situation report 116). These numbers may be a vast underestimation as many of the infected patients may remain asymptomatic and can only be detected by antibody testing^1^. Similarly, many of the deaths may not be accounted due to lack of testing. The disease is caused by a novel coronavirus termed SARS-CoV-2 which belongs to the *Coronaviridae* family and is the third major coronavirus pandemic in last 20 years after Severe Acute Respiratory Syndrome (SARS) and Middle East Respiratory Syndrome (MERS)^2-7^. The lack of available therapeutic options is a major limiting factor in treating these infections which leads to excessive mortality.

Currently, there is an urgent and unmet need of effective antiviral therapy that can not only decrease the disease burden in the patient but can also decrease the ability of the person to infect others. Developing a novel drug many take years to confirm the safety and efficacy, which may not be practical to deal with a widespread pandemic that is killing thousands of people every day. Alternatively, it sounds a lucrative option to repurpose US Food and Drug Administration (FDA) approved drugs for their efficacy against SARS-CoV-2. Earlier screens have found antiviral efficacy of approved therapies such as hydroxychloroquine^8-12^, however, these therapies failed to provide any beneficial effects in COVID-19 due to their toxic side effects^13-18^. Finding efficacy of an approved drug against SARS-CoV-2 with minimal toxicity can provide much needed therapeutic option to treat COVID-19.

Here we screened approximately 1700 US FDA approved compounds to test their ability to inhibit SARS-CoV-2 replication. Our data report finds 24 compounds that are highly effective in inhibiting SARS-CoV-2 replication at concentrations that were significantly lower than those having cytotoxic effects. We also investigated possible mechanism of these compounds.

## Results

### Inhibitory potential of FDA approved drugs against human coronavirus OC43

Initial screening was performed using HCoV-OC43 due to its low biosafety concerns. OC43 is a human coronavirus that usually causes mild disease in humans and cattle^19^. The experimental protocol is demonstrated in Fig. 1A using LLC-MK2 cells which were infected with OC43 at a MOI of 1 for 48 h in presence of the US FDA licensed compounds. The inhibitory potentials of these compounds were measured with the treatment of compounds at 10 μM for 48 hours. The viral presence was detected by immunostaining for the virus and DAPI staining for the cell nuclei. The inhibitory capacity was measured by using the ratio of viral fluorescence to the DAPI and is depicted in Fig. 1B. The initial screen obtained 231 compounds that had ability to inhibit OC43 replication >95%. The remdesivir was used to as a positive control (Fig. 1 C and D).

**Figure 1.**
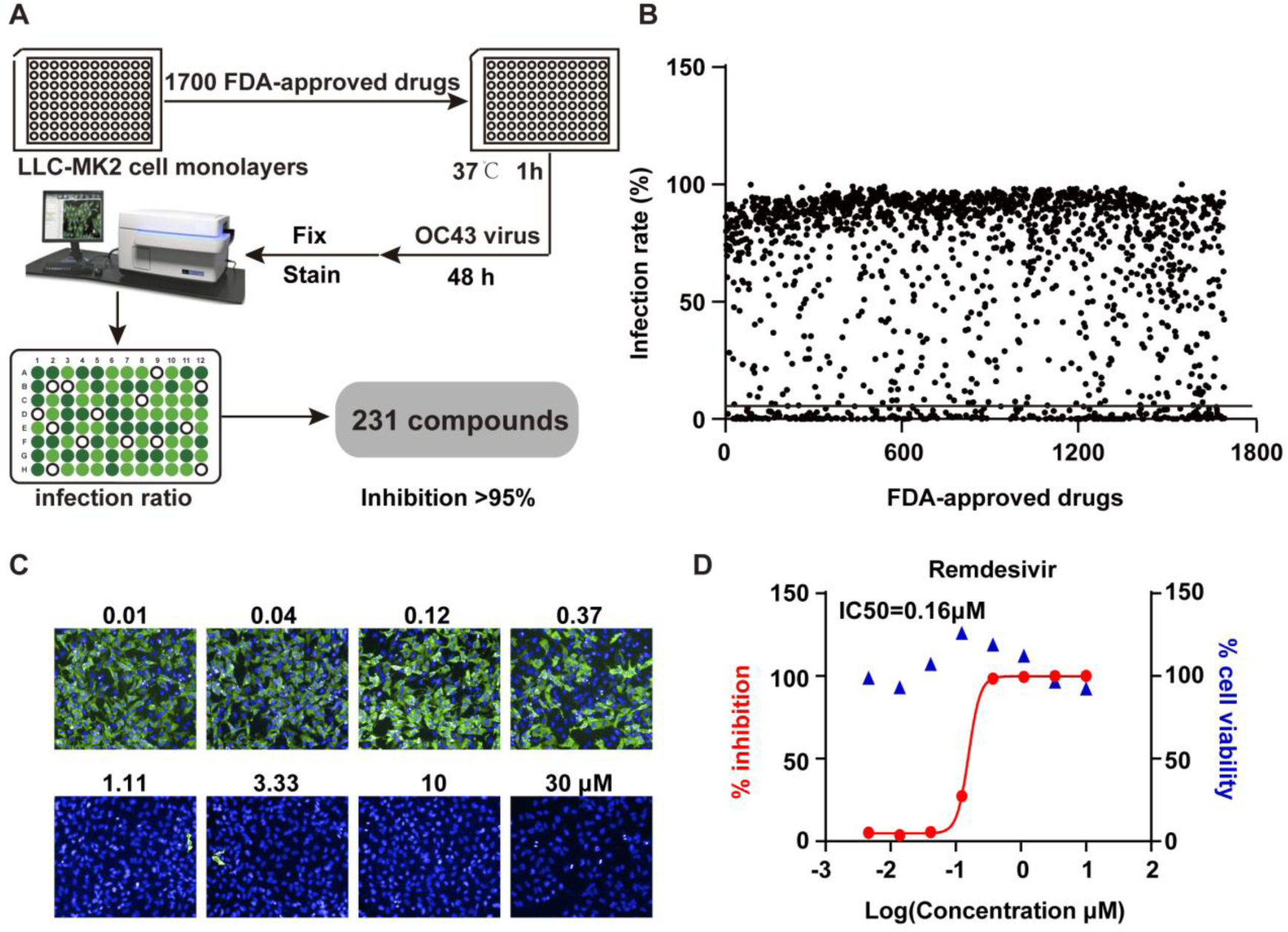
High-throughput screening of US FDA-approved drug library to inhibit human coronavirus OC43 replication *in vitro*. A. The strategy of high throughput screening to identify antiviral drugs that effectively inhibit OC43 replication. LLC-MK2 cells were pretreated with FDA-approved drugs at 10μM for 1h and infected with 1 MOI of OC43 for 48h. Cells were then fixed and stained to calculate the infection ratio with Operetta software. B. Primary screening results of 1700 FDA-approved drugs against OC43, each dot represents one compound along with the rate of OC43 inhibition. C. Image samples show signals of OC43 infection in cell cultures. LLC-MK2 cells were treated with indicated doses of remdesivir for 1h, and then cells were infected with OC43 for 48 h. D. Cell viability of remdesivir to LLC-MK2 cells were measured by CCK-8 assays. The % inhibitions were calculated according to the data in C.

### Calculation of IC50, CC50 and SI of FDA approved drugs against OC43

Next, we sought to determine the effective concentrations of positively screened drugs in our initial approach and test the toxicity profile of these drugs in relation to their viral inhibitory concentrations. Our data show that vast majority of the positively screened drugs were effective in inhibiting the viral replication at sub-micromolar concentrations including many of them can almost completely inhibit the viral replication at micromolar range (Fig. 2). Surprisingly, the effective drugs against the coronavirus belonged to a wide range of therapeutic groups including those used for neurological diseases, hormones, enzymes and antimicrobial agents among others (Table 1). The cytotoxic concentration to kill 50% cells (CC50) was noted for these drugs by measuring cell viability over similar concentrations. The selective index in our study was found to be >600 for 5 of the screened compounds. The SI for the hydroxychloroquine was 22 in our study, indicating increased safety of newly identified drugs compared to the hydroxychloroquine.

**Table. 1.**
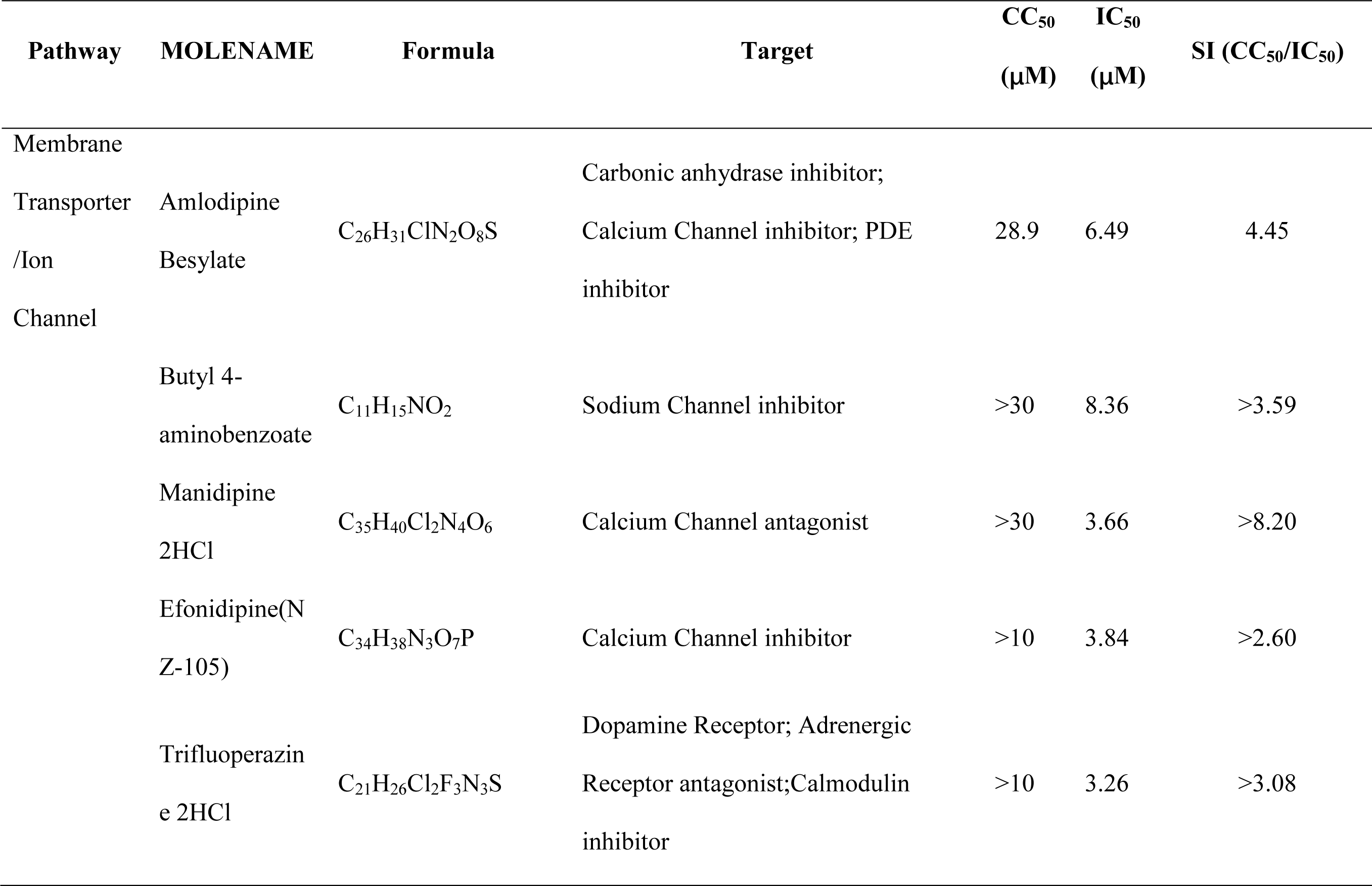
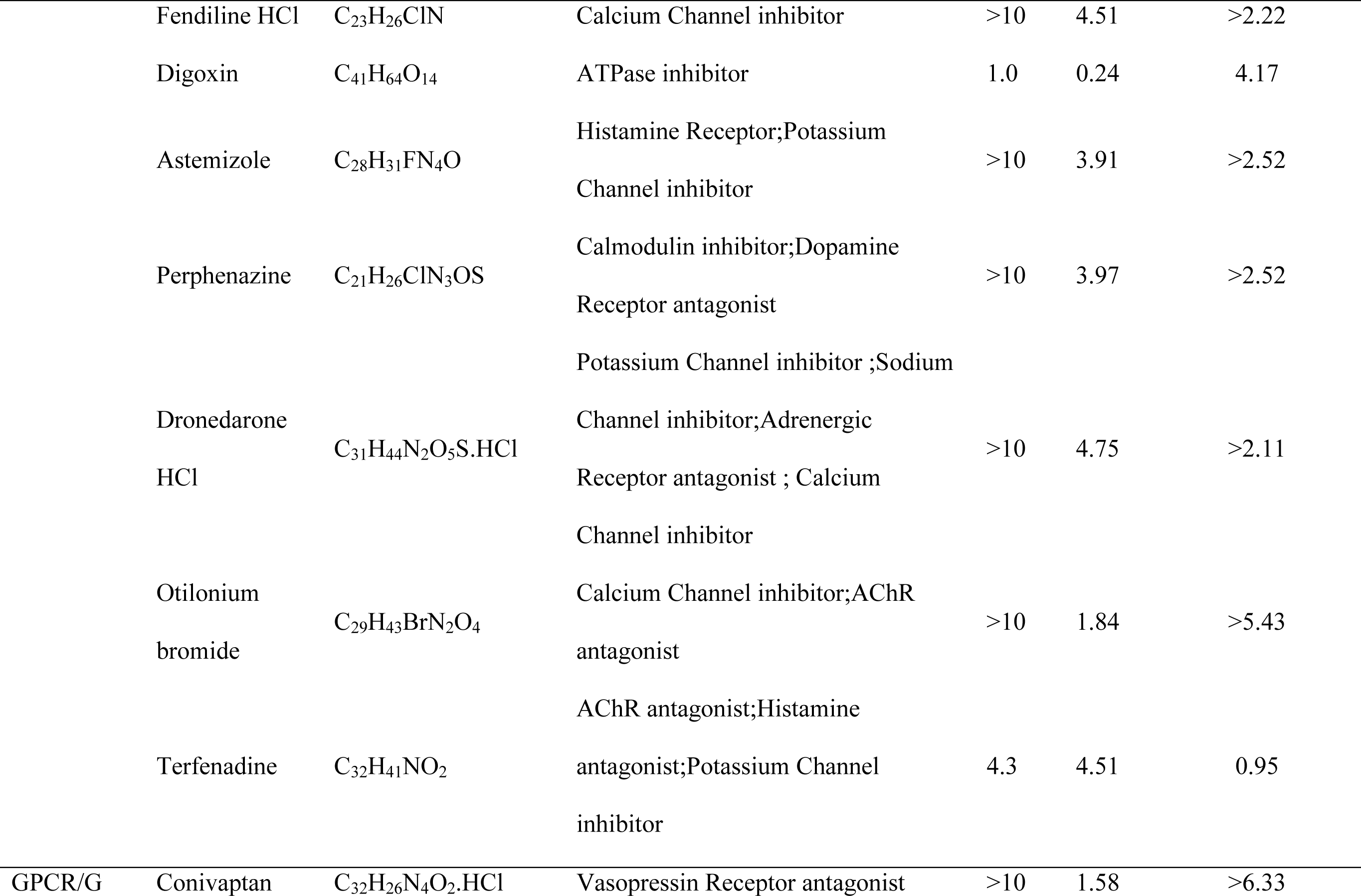
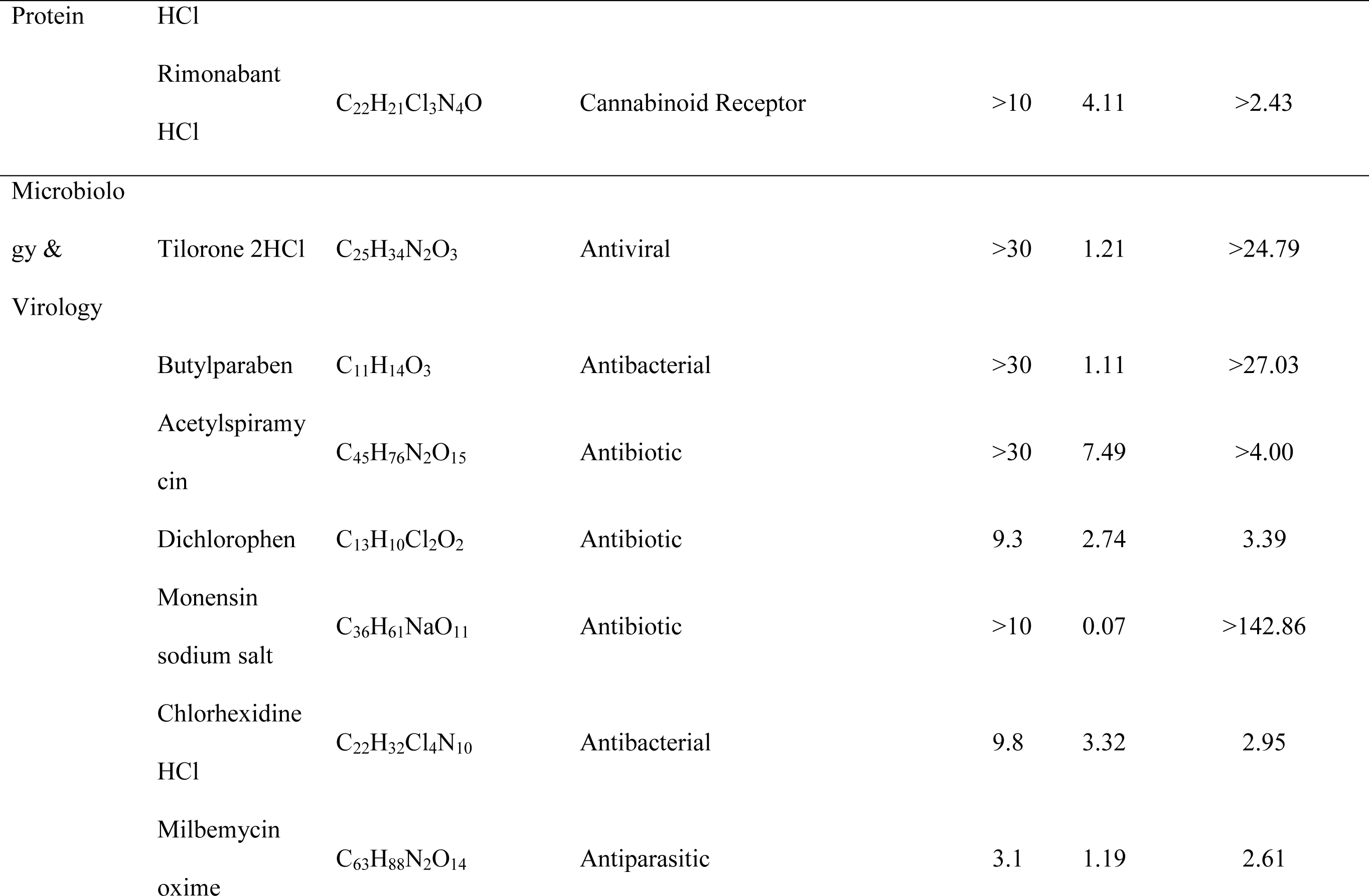
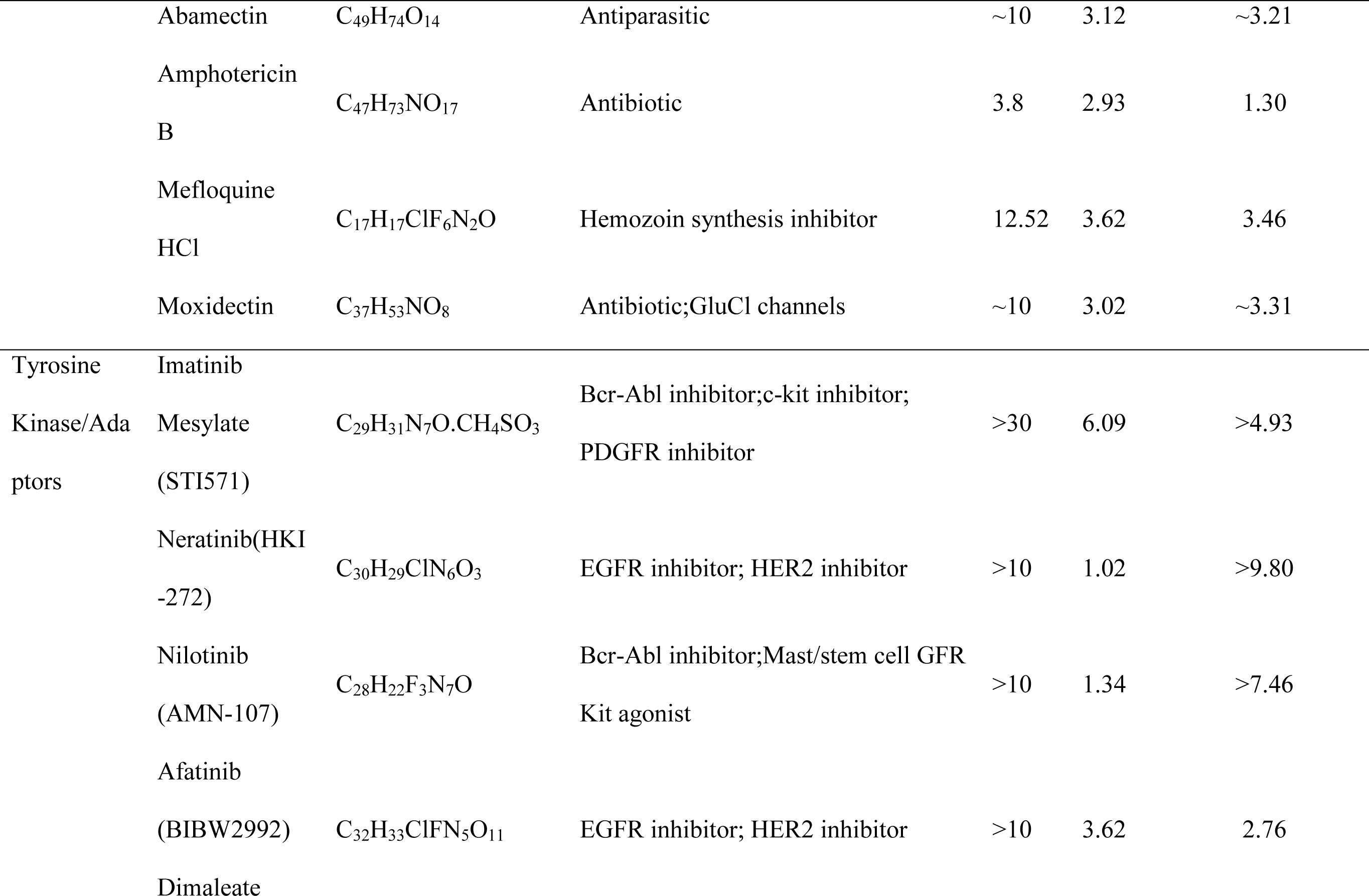
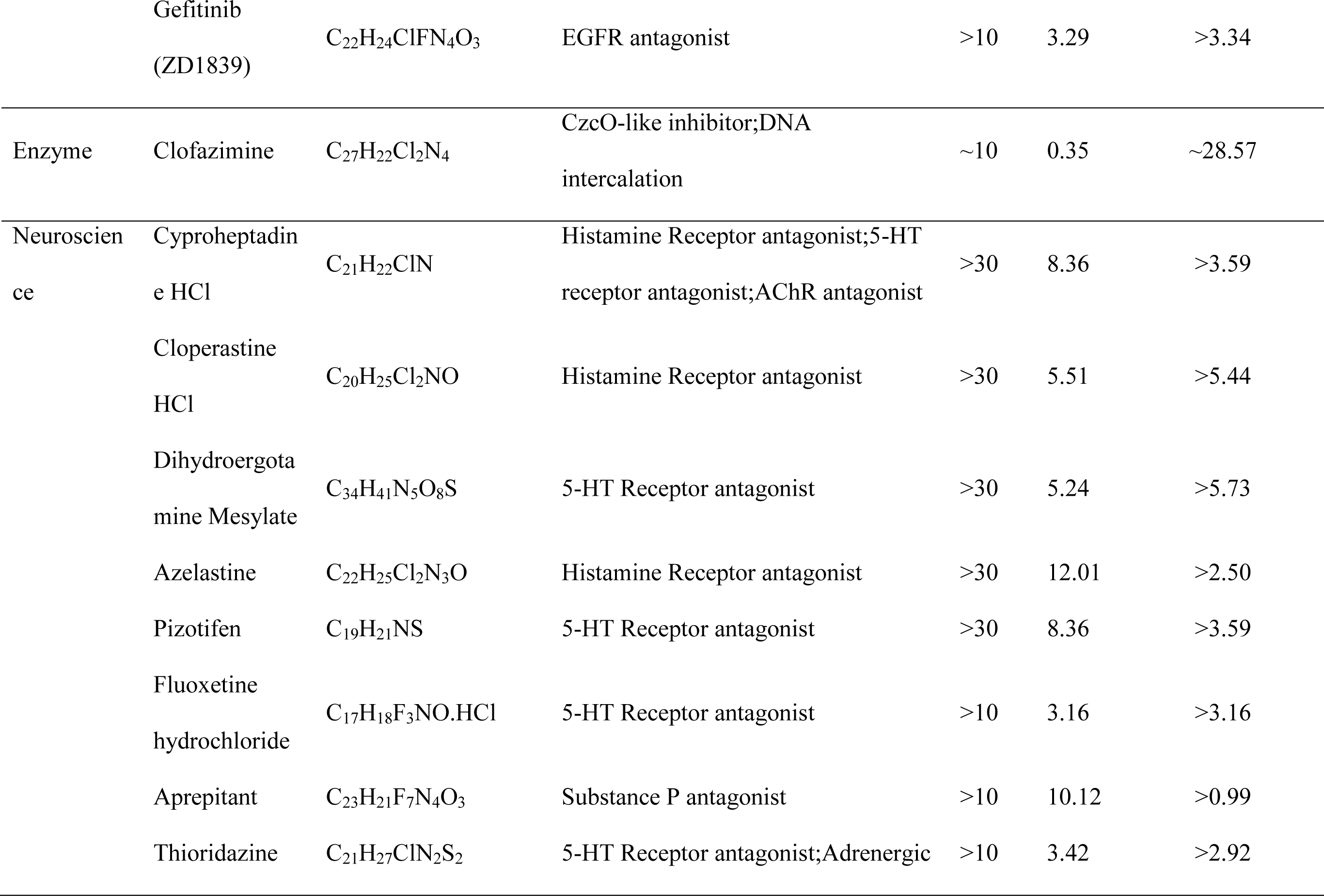
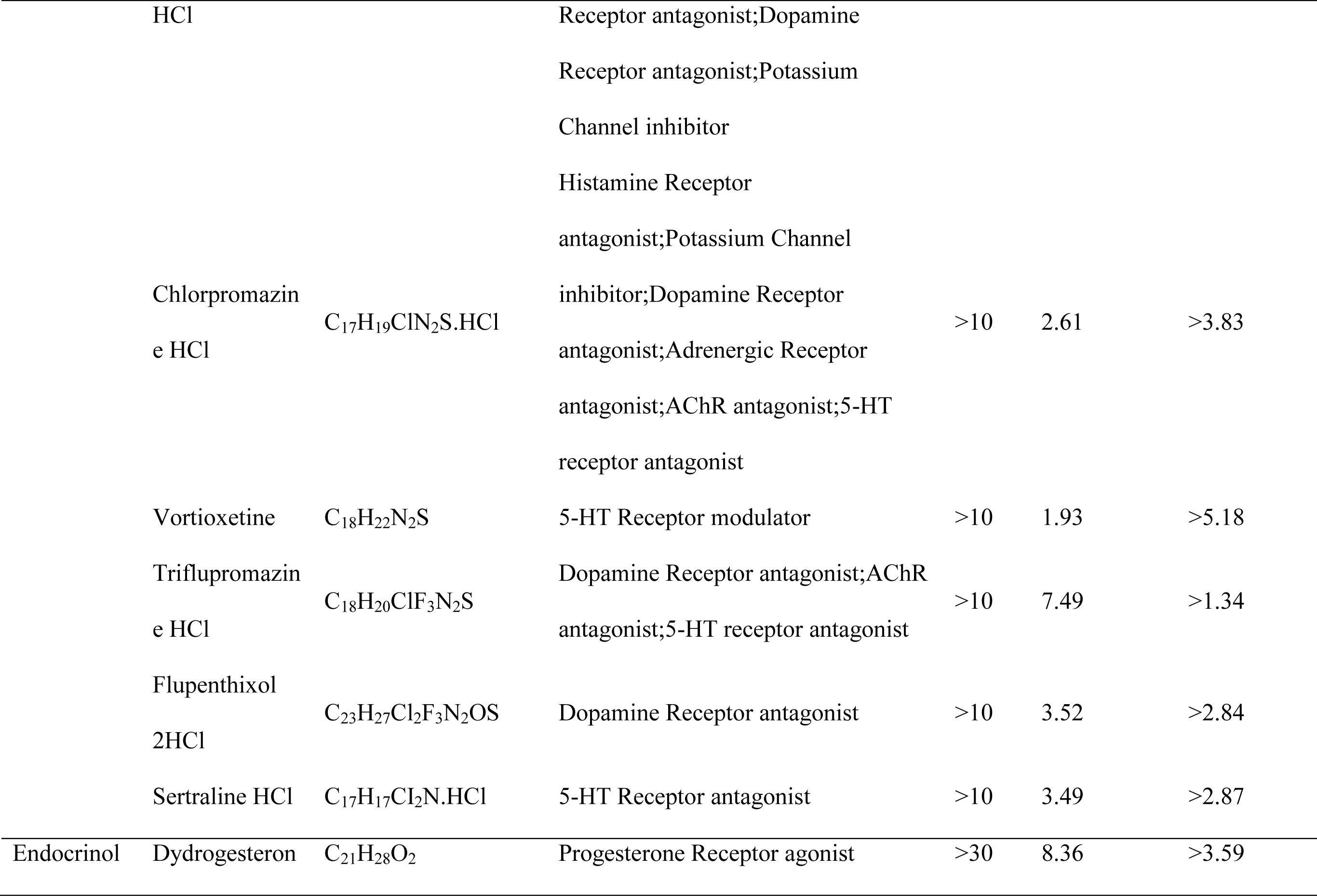
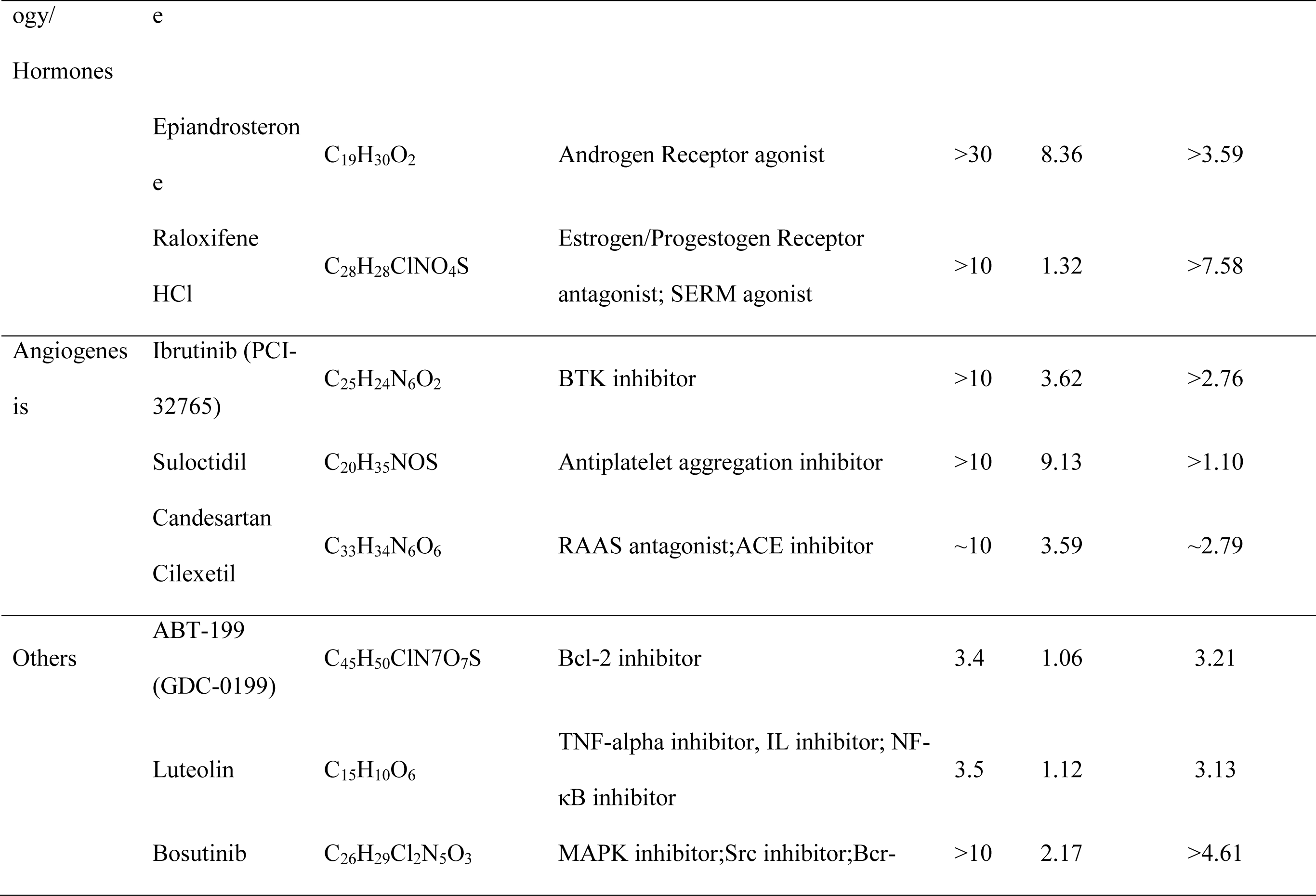
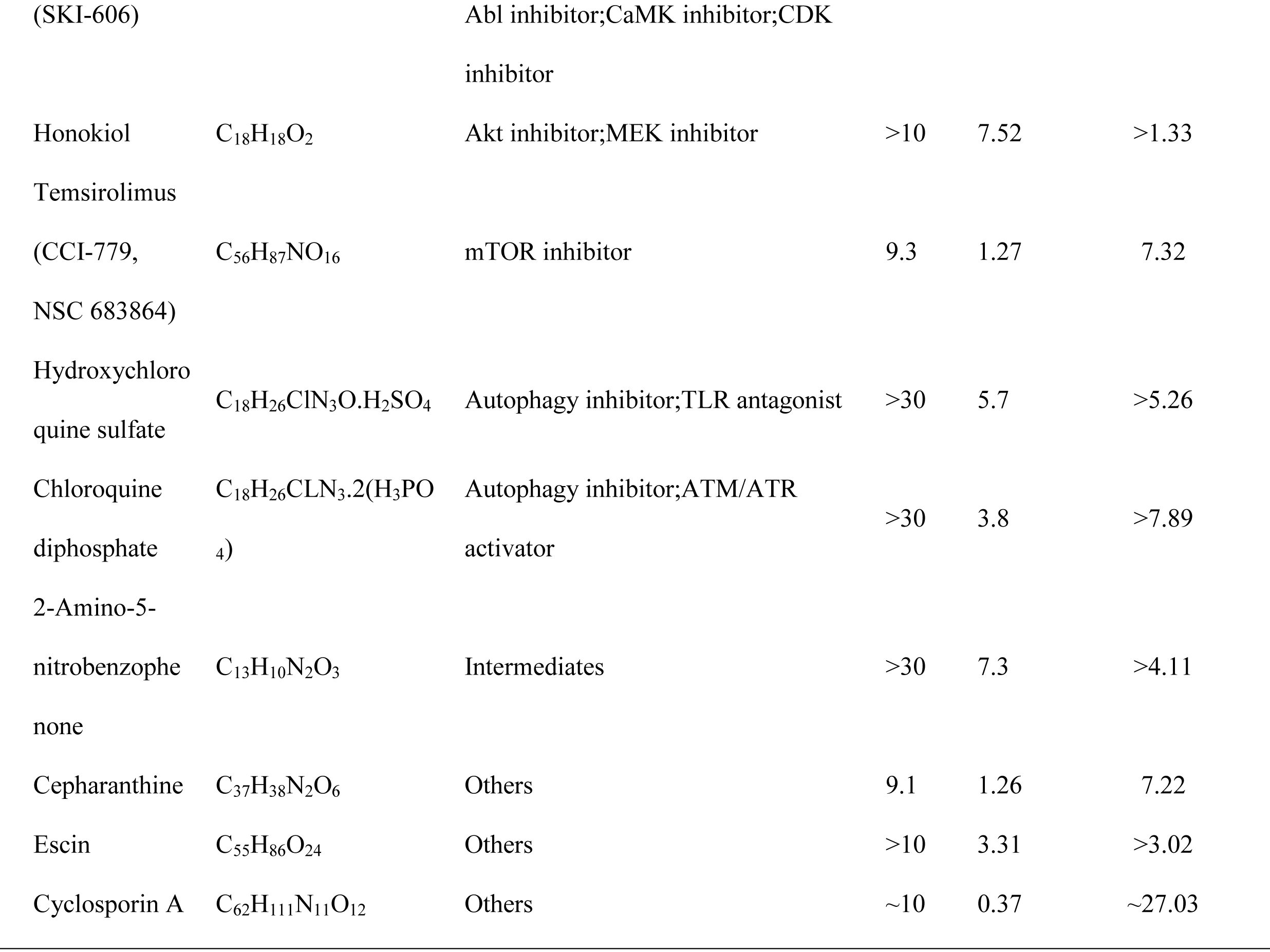
Antiviral activity of selected compounds against OC43.

**Figure 2.**
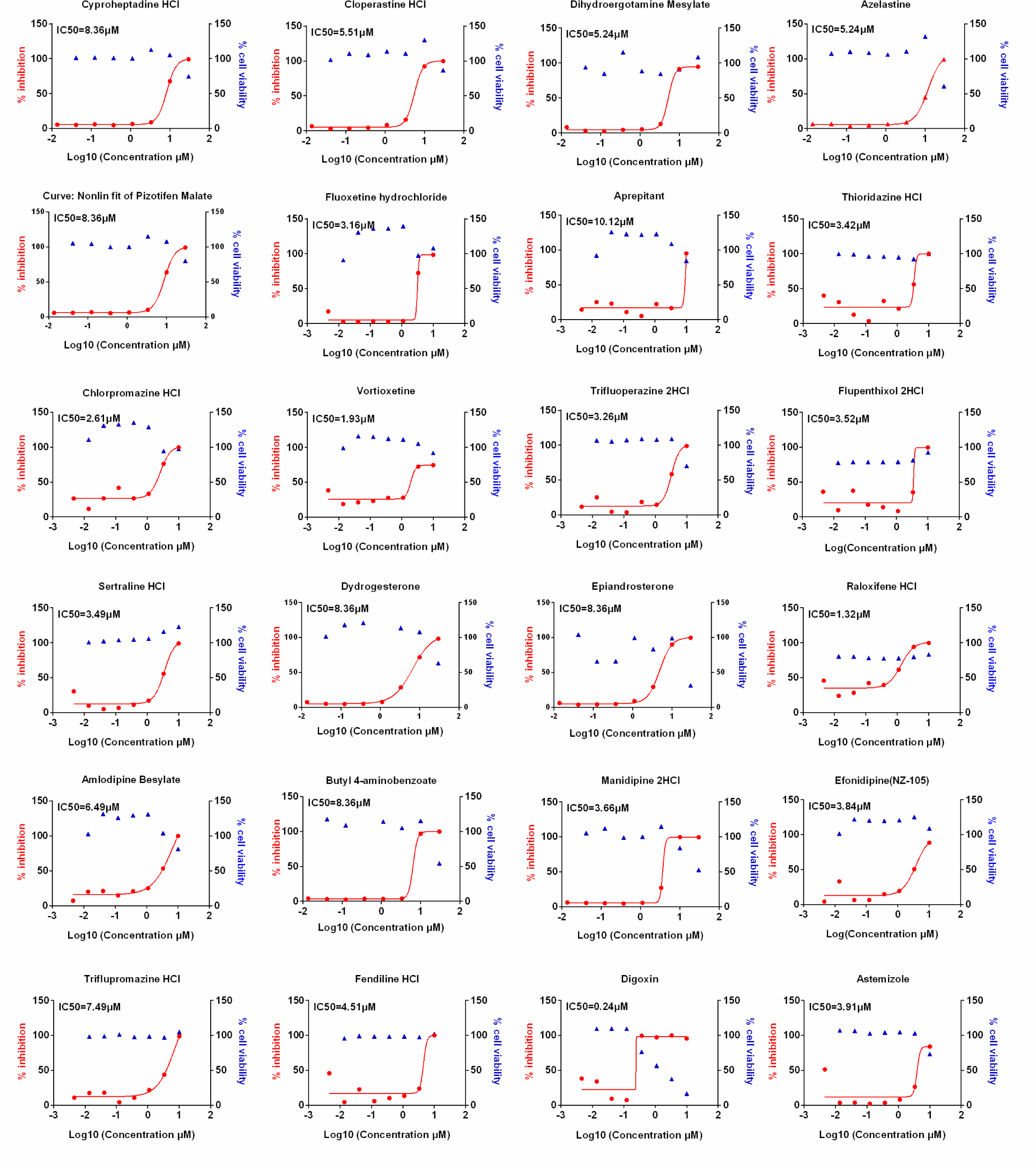
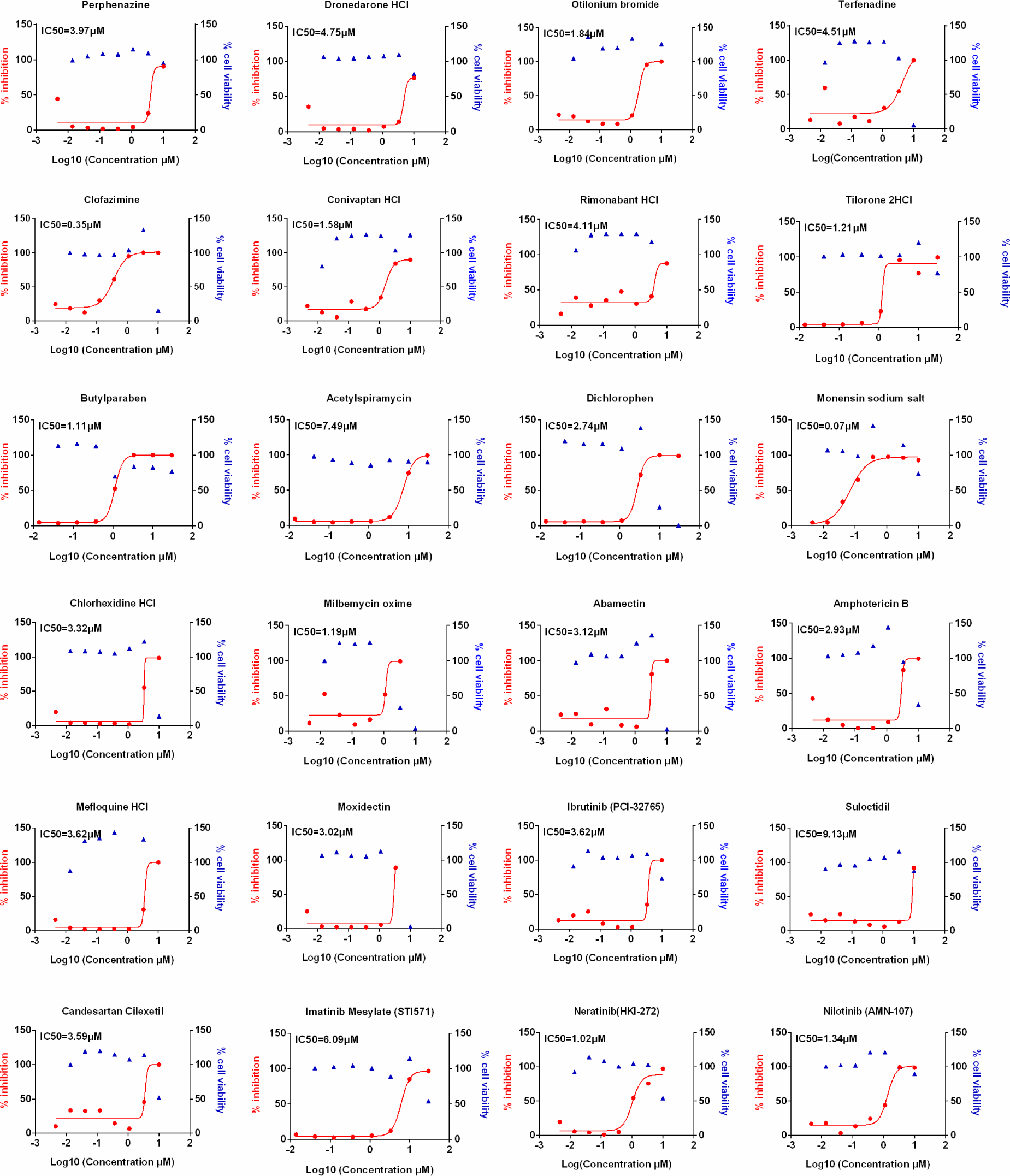
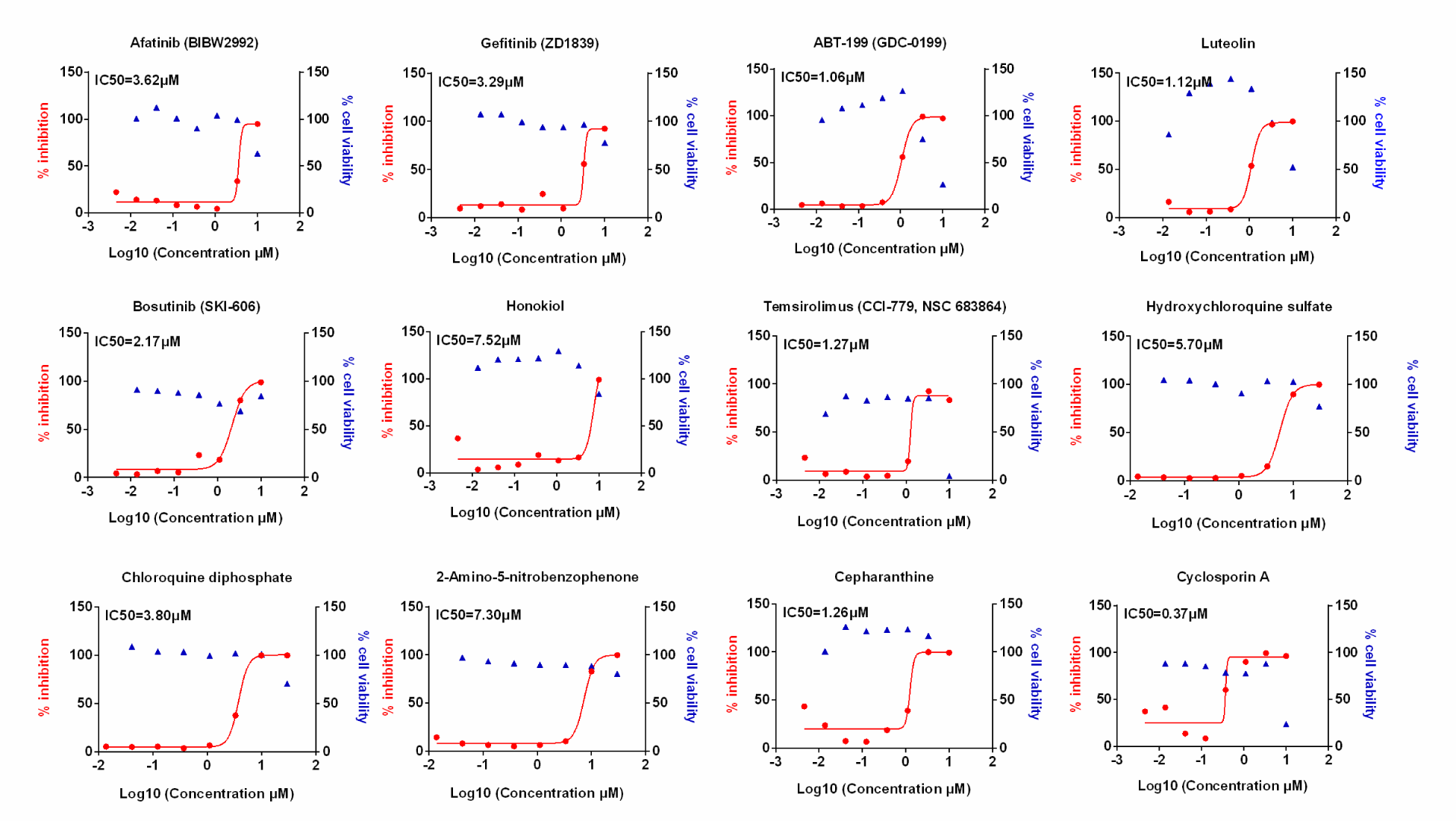
Dose response curves of selected compounds from the hits against OC43 infection *in vitro*. LLC-MK2 cells were pretreated with indicated drugs at 37*C for 1h with eight doses (0.014, 0.041, 0.123, 0.370, 1.111, 3.333, 10, 30μM) with three-fold dilution followed by infection with OC43 at MOI of 1 for 48h. In parallel, effects of these compounds on the cell viability in LLC-MK2 cells were measured by CCK-8 assays. The left *Y*-axis of the graphs represent % inhibition of the infection and the right *Y*-axis of the graphs present % cell viability in presence of the drugs.

### Testing the antiviral effectiveness of positive drugs against SARS-CoV-2

Finally, we sought to determine whether these compounds can show efficacy against SARS-CoV-2, the causative agent of COVID-19. Positive compounds from the initial screen were tested for their antiviral efficacy against SARS-CoV-2 in Vero cells. SARS-CoV-2 replicates within the Vero cells and causes cytopathic effects in these cells in absence of any antiviral treatment. We generated the dose response inhibition curves along with the cytotoxicity curves for these compounds in presence of SARS-CoV-2 (Fig. 3). Our data show that 24 of these compounds show significant efficacy in inhibiting SARS-CoV-2 replication with sub micromolar IC50 for many of these drugs such as nilotinib, clofazimine and raloxifene. The effects also confirmed by immunofluorescence assay (data not shown). These compounds also belong to a wide variety of classes including cardiac glycosides, anti-malarial drug hydroxychloroquine, cyclooxygenase-2 inhibitors and ion channel blockers, among others. The IC50, CC50 and SI of these compounds is shown in Table 2.

**Table. 2.**
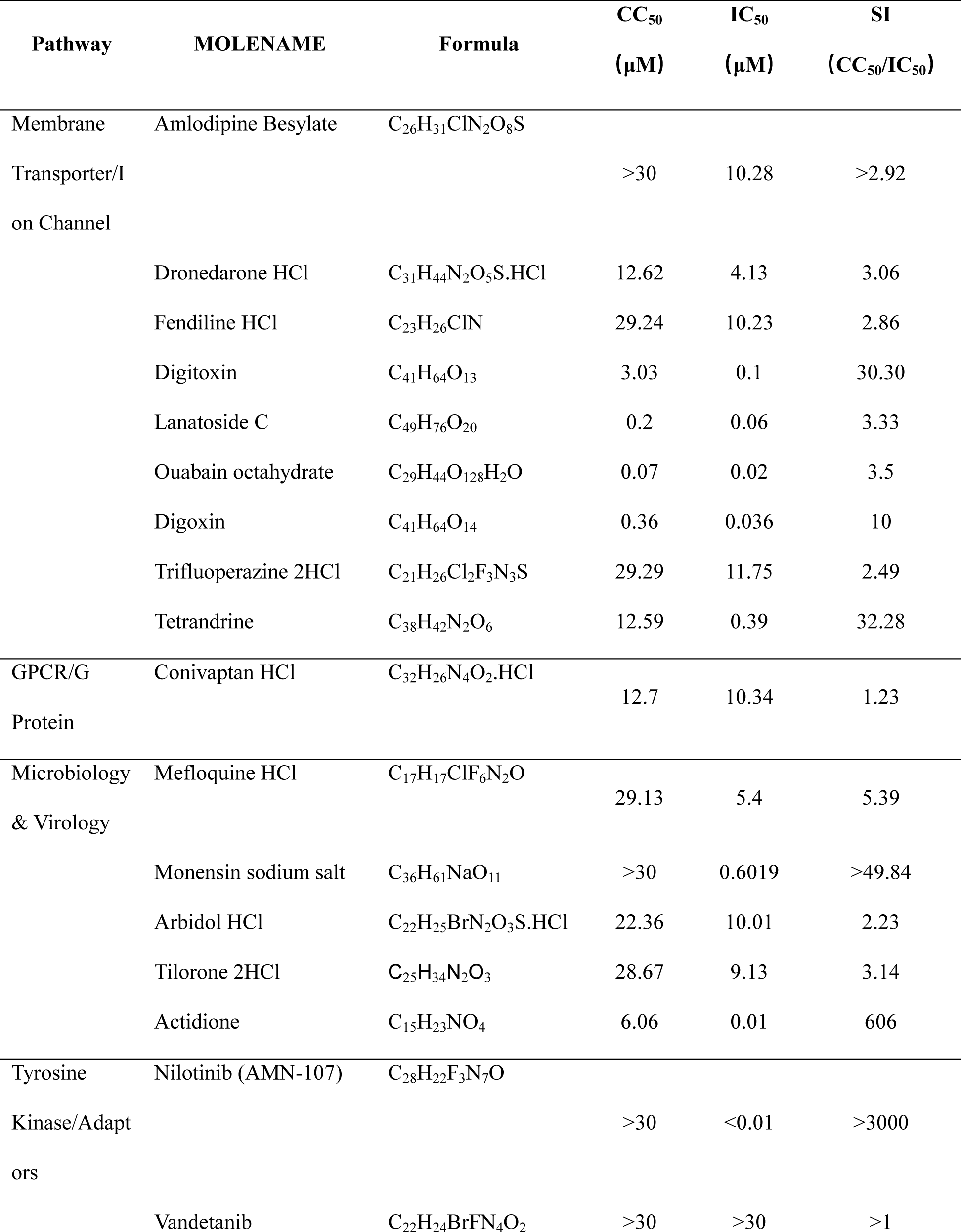
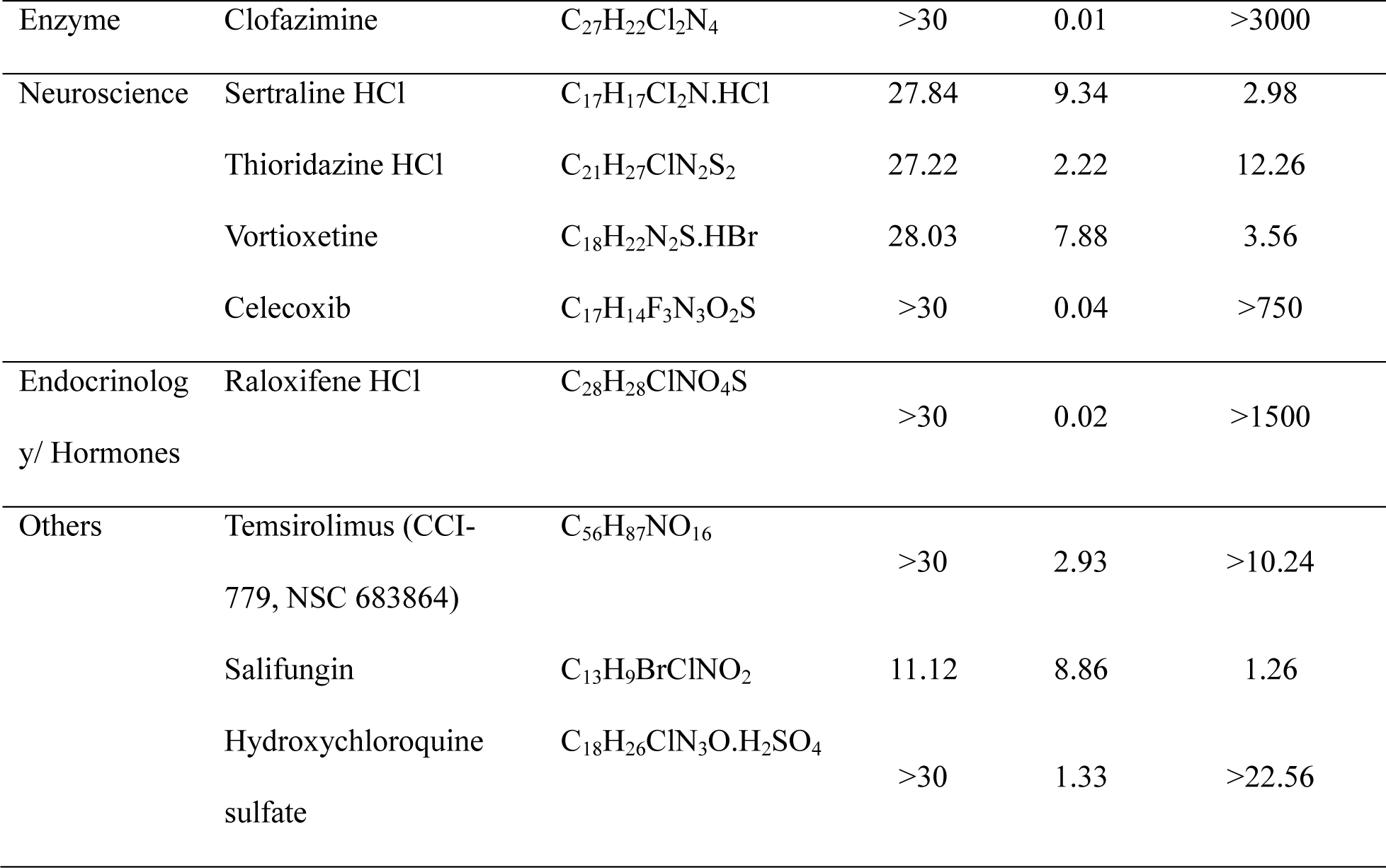
Antiviral activity of selected compounds against SARS-CoV-2.

**Figure 3.**
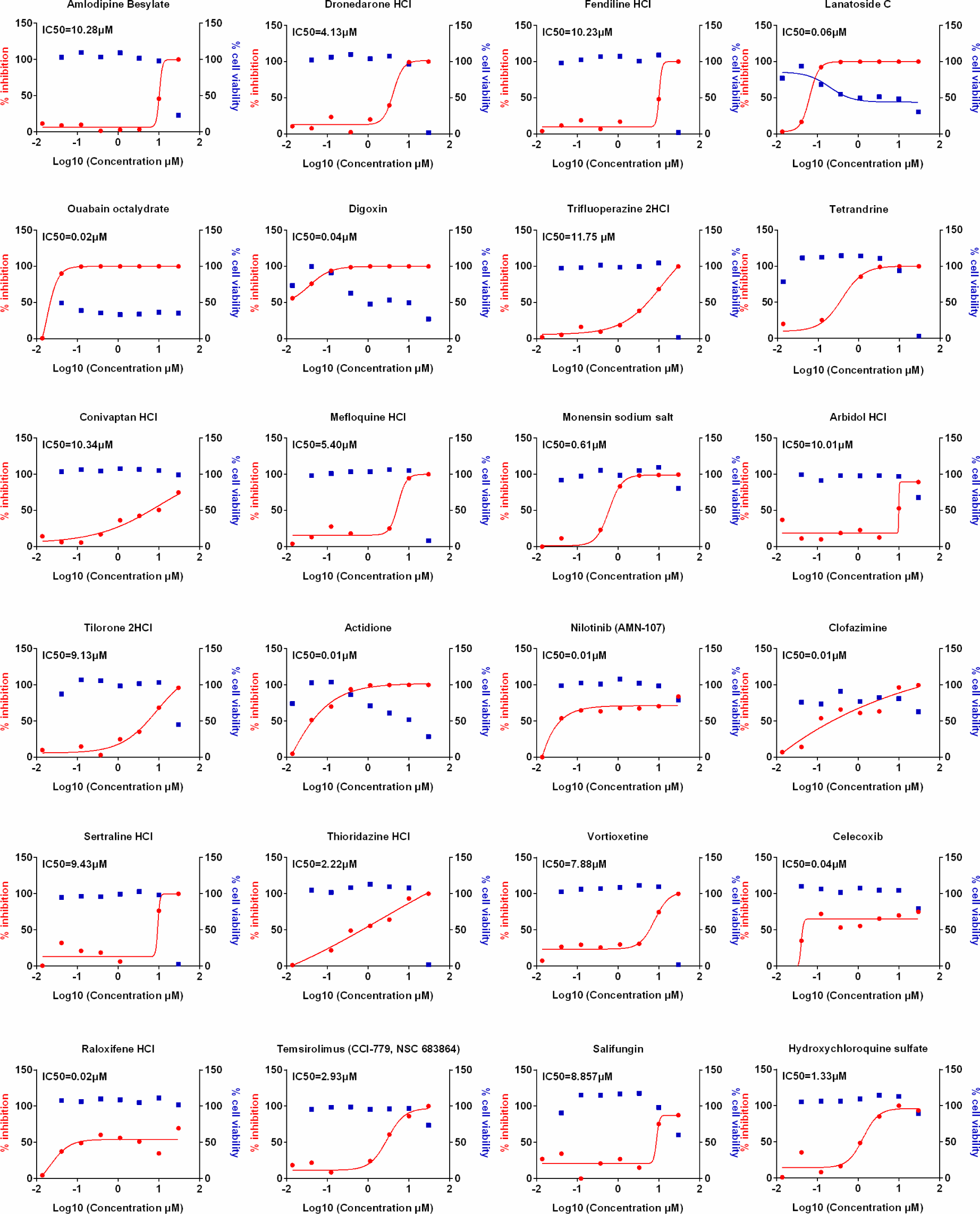
Dose response curves of selected compounds against SARS-CoV-2 infection. Vero cells were pretreated with indicated drugs at 37*C for 1h with eight doses (0.014, 0.041, 0.123, 0.370, 1.111, 3.333, 10, 30μM) with three-fold dilutions, followed by infection with SARS-CoV-2 at a MOI of 0.1 for 24h. The viral load in the cell supernatant was quantified by qRT-PCR. Meanwhile, cell viability in presence of these drugs were measured in Vero cells by CCK-8 assays. The left *Y*-axis of the graphs represent % inhibition of viral replication and the right *Y*-axis of the graphs indicates % cell viability in presence of the drugs.

### Five candidate drugs inhibit cell fusion

Finally, in order to clarify the mechanism by which the compounds inhibit SARS-CoV-2, we investigated the effects of these candidates on virus entry. First, we constructed the cell-cell fusion assays. As indicated in Fig 4, SARS-CoV-2 S protein expression can initiate cell fusion with ACE2-overexpressed cells, but the control vector did not. Then, we detected the effects of these indicated drugs on S protein-mediated cell fusion. Indicated drugs were added to cells before co-incubation of the cells. We found that digoxin and ouabain octahydrate induced cell death, but not other indicated drugs. Interestingly, fendiline hydrochloride, monensin sodium salt, vortioxetine, sertraline hydrochloride, and raloxifene hydrochloride inhibited the SARS-CoV-2 S protein-mediated cell fusion (Fig. 4).

**Figure 4.**
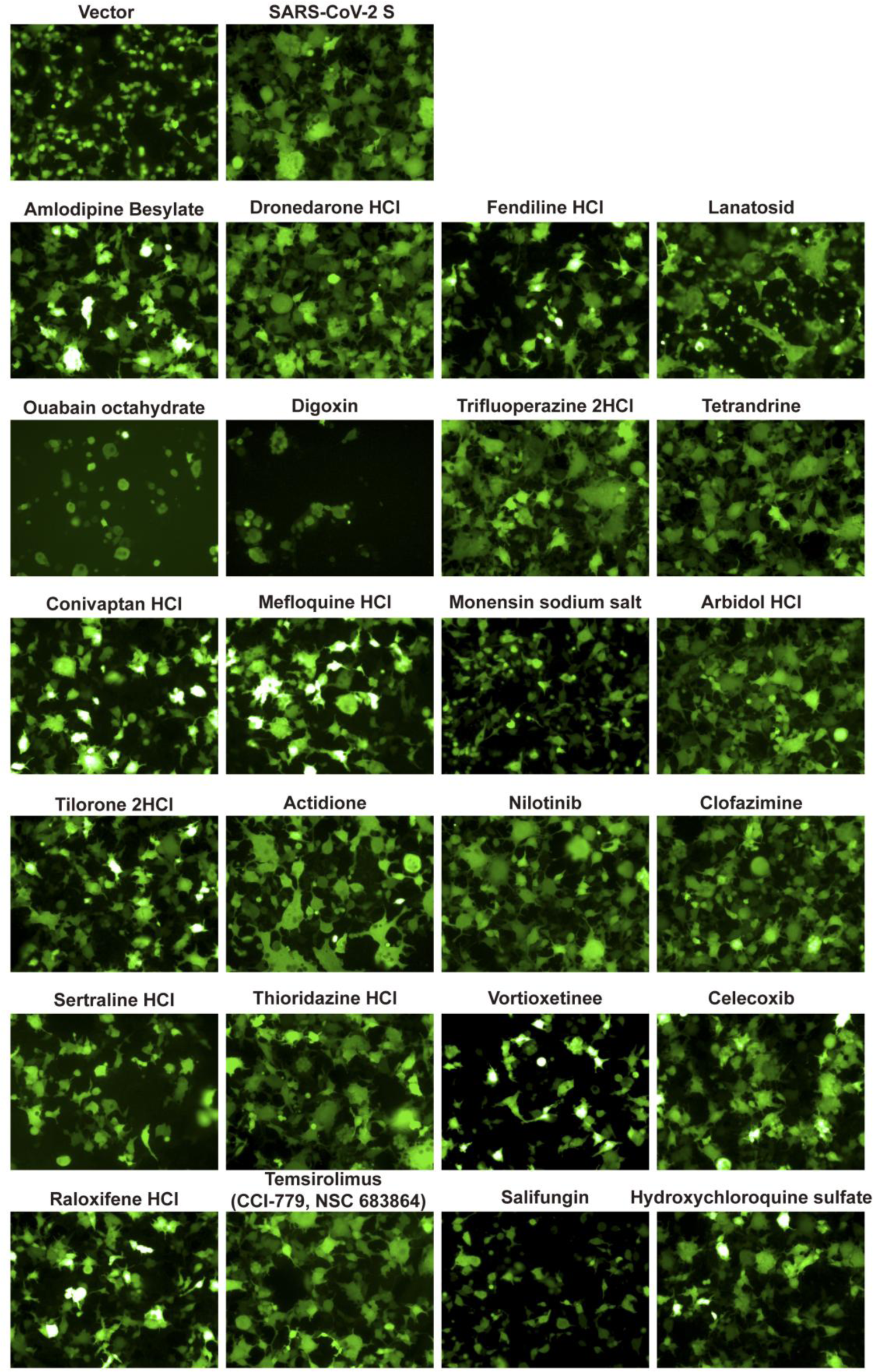
The effect of indicated drugs on cell-cell fusion mediated by SARS-CoV-2 S protein. HEK-293T cells were co-transfected with SARS-CoV-2-S glycoprotein and eGFP. 24 h post transfection, cells were digested with trypsin (0.25%) and overlaid on a 50% confluent monolayer of 293T-ACE2 cells at a ratio of 1:1 which were treated with candidate drugs for 1h. After 24 h incubation, images of syncytia were captured with Operetta and analyzed by Harmony software.

## Discussion

The current pandemic of COVID-19 is the third major outbreak in this century and the largest outbreak of the coronavirus in known human history. The three novel coronavirus outbreaks in such a short span of time are strong indicators of potential threat posed by coronaviruses. While majority of the respiratory viral infection research has been focused on influenzas viruses which causes huge burden of seasonal flu and occasionally pandemic outbreaks, coronavirus is likely to emerge a similar or more severe pathogen than flu in long term.

Given the scale and devastation of the current COVID-19 outbreak, and persistent threat of coronaviruses in causing human disease, there is urgent need to find effective and safe therapies that can treat these patients. Currently, there are no approved therapies for coronaviruses including SARS-CoV-2. The experimental therapies being used with known antiviral agents either show limited efficacy (remdesivir) or have high systemic toxicity (hydroxychroquine), limiting their usefulness^20-22^. Finding new therapies that are effective and safe are urgently needed. In this study, we have identified many FDA approved therapies that are highly effective against coronaviruses, including 24 of the agents that are effective against SARS-CoV-2. This screening confirms previous reports demonstrating anti-SARS-CoV-2 activity of hydroxychroquine, amlodipine besylate, arbidol hydrochloride, tilorone 2HCl, dronedarone hydrochloride, tetrandrine, merfloquine, and thioridazine hydrochloride^8-10, 23^, while identifying additional 14 drugs. The underlying mechanisms of viral replication inhibition by these drugs is not clear. It is highly unlikely that these compounds will have similar antiviral mechanisms given the vast structural and pharmacological diversities of the effective antiviral compounds in our study. However, it is clear from other viral studies such as influenza or HIV, where antiviral drugs can affect various steps in the viral life cycle including attachment, entry, replication, assembly and budding of viral progeny. Five drugs may inhibit S-mediated cell fusion as indicated by our data (Fig. 4). Further studies are required to understand the precise mechanisms of each of the effective compounds found in this study.

Toxicity is one of the limiting factors in the therapeutic application of many drugs despite their known antiviral activities. Many of these drugs had SI of > 600, showing promise of their usefulness at safe doses. For comparison, the SI of hydroxychloroquine was found to be 22 in our study while SI of amlodipine besylate was found to be ∼3, demonstrating much lower safety profile of this drug. Similarly, other drugs that are known to have low selective index such as digoxin for their approved use, also show lower SI in our screen. Five of the drugs with SI of >600 include tyrosine kinase inhibitor nilotinib, antibiotics such as clofazimine and actidione, selective estrogen receptor modulator such as raloxifene and non-steroidal anti-inflammatory drug celecoxib.

etacoronaviruses have raised great public health threats to human beings, as most known HCoVs including all the three virulent HCoVs (SARS-CoV, MERS-CoV and SARS-CoV-2) and two seasonal HCoVs (OC43 and HKU1) belong to this species^3-7, 19, 24^. It is of great value to identify antivirals against a broad spectrum of HCoVs, particularly the Betacoronaviruses, to tackle such threats by pharmaceutical interventions. To this end, we first screened the compounds which showed apparent activity of anti-OC43, the most prevalent HCoV circulates worldwide^25^. We then narrowed down the candidates by the screening on SARS-CoV-2, resulting in the identification of 24 compounds which can inhibit both OC43 and SARS-CoV-2. Our study provides a foundation for subsequent anti-HCoVs drug screening of broad spectrum. However, further tests are warranted to verify their efficacies.

In summary, our screen identified 14 previously unknown FDA approved compounds that are effective in inhibiting SARS-CoV-2 beside confirming the antiviral properties of ten previously reported compounds, validating our approach. This screen identified five new compounds that are highly efficacious in inhibiting the viral replication of SARS-CoV-2 with SI >600. Further studies are needed to confirm the *in vivo* efficacy of these drugs in humans and COVID-19 relevant mouse models such as those with human ACE2 transgene^26^.

## Materials and Methods

### Cells and Viruses

LLC-MK2 cells (Rhesus monkey kidney cells) were cultured in 64% Hank’s MEM and 32% Earle’s MEM (Gibco, New York, USA) supplemented with 3% fetal bovine serum (FBS) (Hyclone, Utah, USA) and 1% glutamine (Thermo, Massachusetts, USA). Vero cells (African green monkey kidney cell) were cultured in Dulbecco’s Modified Eagle’s Medium (DMEM, Gibco) supplemented with 10% FBS.

Human coronavirus (HCoV) strain OC43 was propagated in LLC-MK2 cells in 0.5% FBS MEM and virus titers were determined via TCID50 with LLC-MK2 cells. SARS-CoV-2 virus was isolated from the respiratory samples of patients in Wuhan of Hubei Province *(3)*. SARS-CoV-2 virus was propagated in Vero cells and used in this study. All experiments with SARS-CoV-2 virus were conducted in BSL-3 laboratory.

HEK293T cells stably expressing recombinant human ACE2 (293T/hACE2) were maintained in Dulbecco’s MEM containing 10% fetal bovine serum and 100 unit penicillin, and 100μg streptomycin per milliliter.

### Antibodies

Mouse polyclonal against OC43 N antibody was prepared in the laboratory. Rabbit polyclonal against SARS-CoV-2 N protein antibody was purchased from Sino Biological (Beijing, China). Alexa Fluor 488-conjugated goat anti-mouse IgG, Alexa Fluor 488-conjugated goat anti-rabbit IgG were purchased from Thermo.

### Screening of FDA-approved drugs

US FDA-approved drug library which contain 1700 compounds was purchased from TargetMol (Massachusetts, USA). LLC-MK2 cells were seeded at 2×10 ^4^ cells per well in 96-well plates and incubated at 37°C and 5% CO_2_. The next day, LLC-MK2 cells were treated with the compounds at a concentration of 10μM. After 1 h of treatment, cells were infected with OC43 at MOI of 1. At 48 h post infection (hpi), cells were fixed with 4% paraformaldehyde for 20 min at room temperature. Immunofluorescence staining was performed using mouse anti-OC43 NP antibody, followed by anti-mouse Alexa Flour 488 and DAPI (Sigma, St. Louis, MO). Images were captured by Operetta (PerkinElmer, Massachusetts, USA) at the magnification of 20× objective. The infection ratios were calculated using automated image analysis software (Harmony 3.5.2, PerkinElmer). Remdesivir and DMSO were used as positive and negative controls, respectively.

The positively identified drugs from this screen were used to perform dose response curves against OC43 on LLC-MK2 and against SARS-CoV-2 using Vero cells as described below.

### IC_50_ (The half maximal Inhibitory concentration), CC_50_ (The half maximal cytotoxic concentration) and SI (Selectivity index) determination

LLC-MK2 cells (for OC43 infection) or Vero cells (for SARS-CoV-2 infection) were seeded in 96 wells plate one day before infection at the concentration of 2×10^4^ cells /well or 1.4×10^4^ cells /well, respectively. For IC50, cells were pre-treated for 1h with each drug at concentrations 0.013, 0.041, 0.123, 0.370, 1.111, 3.333, 10, 30 μM, and then infected with virus at MOI of 1. At 48 hpi (OC43) or 24 hpi (SARS-CoV-2), cells were fixed with 4% paraformaldehyde for 20 min at room temperature. Immunofluorescence was conducted with mouse anti-OC43 N protein antibody, or rabbit anti-SARS-CoV-2-NP antibody, and followed by anti-mouse, or anti-rabbit Alexa Flour 488 and DAPI. Images were performed by Operetta with 20× objective. The IC _50_ were calculated using automated image analysis software (Harmony 3.5.2, PerkinElmer).

For CC_50_, cells were pre-treated with each drug at concentrations 0.013, 0.041, 0.123, 0.370, 1.111, 3.333, 10, 30 μM, respectively. After 48h (LLC-MK2 cells) or 24h (Vero cells) post treatment, cell viability was evaluated by using a CCK8 kit (Yeasen, Beijing, China) according to the manufacturer’s instructions. Selectivity index was calculated using following formula: SI=CC_50_/IC_50._ Graphpad Prism 7.0 was used for analyzing IC_50_ and CC_50_.

#### Immunofluorescence

Cells were fixed with 4% paraformaldehyde for 20 min at room temperature, and permeabilized with 0.5% Triton X-100 for 10 min. Cells were then blocked with 5% BSA and stained with primary antibodies, followed by staining with an Alexa Fluor 488 secondary antibodies. Nuclei were counterstained with DAPI.

### Quantitative RT-PCR

Vero cells were pre-treated with indicated concentrations of drugs for 1h and incubated with SARS-CoV-2 at 0.1 MOI for 1h. Then, cells were washed with opti-MEM for one time and incubated with indicated concentrations of drugs. At 24 hpi, supernatants were collected and viral RNA in the cell supernatants were extracted by using Direct-zol RNA MiniPrep kit (Zymo research, CA, USA) according to the manufacturer’s instructions. Viral copy numbers were measured by RT-PCR using primers and probe targeting the SARS-CoV-2 N gene. The reference standard was tenfold diluted from 1×10^9^ copies to 1×10^4^ copies. PCR amplification procedure was 50°C, 15min, 95°C, 3min; 95°C, 15s, 60°C, 45s+Plate Read,50 cycles. The amplification process, fluorescence signal detection, data storage and analysis were all completed by fluorescence quantitative PCR and its own software (Bio-Rad CFX Manager). The copies of virus were calculated according to the standard curve. The inhibition ratio was obtained by dividing the number of copies of the virus in the vehicle control group. The data were nonlinear fitting by graphpad 7.0 software to calculate IC_50_ of each drug.

### Cell-cell fusion assay

Cell-cell fusion assays were performed as described previously *(19)*. Briefly, HEK-293T cells were co-transfected with SARS-CoV-2-S glycoprotein and eGFP. At 24 h post transfection, cells were digested with trypsin (0.25%) and overlaid on a 50% confluent monolayer of 293T-ACE2 cells at a ratio of 1:1 which were treated with candidate drugs for 1h. After 24h incubation, images of syncytia were captured with Operetta (PerkinElmer, Massachusetts, USA).

## Acknowledgments

This work was supported by grants from the National Major Sciences & Technology Project for Control and Prevention of Major Infectious Diseases in China (2018ZX10301401 to XL), the National Natural Science Foundation of China (81930063, 81971948 to JW and XL), Chinese Academy of Medical Sciences (CAMS) Innovation Fund for Medical Sciences (2016-I2M-1-014, 2016-I2M-1-005 to JW and XL), National Key R&D Program of China (2020YFA0707600 to XL).

## Author contributions

Project conception: J.W., X.L., and L.S.; Experimental design: J.W. X.L., D.C., L.S., Z.Z., and L.R.; Experimental work: X.X., C.W., Y.W., X.D., T.J., X.X., C.W., Z.T., Y.W.; Data analysis: J.W., X.L., L.R., C.S.D.C., X.X., and Z.Z., Writing Original Draft: J.W., X.L., L.S., D.C., and X.X.; Writing Review & Editing, J.W., X.L., D.C., L.S. and C.S.D.C.; All authors reviewed the manuscript.

## References

1. Guo, L, Ren, L, Yang, S, Xiao, M, Chang Yang, F, et al. (2020). Profiling Early Humoral Response to Diagnose Novel Coronavirus Disease (COVID-19). Clinical infectious diseases : an official publication of the Infectious Diseases Society of America.

2. Huang, C, Wang, Y, Li, X, Ren, L, Zhao, J, Hu, Y, et al. (2020). Clinical features of patients infected with 2019 novel coronavirus in Wuhan, China. Lancet 395: 497–506.

3. Ren, LL, Wang, YM, Wu, ZQ, Xiang, ZC, Guo, L, Xu, T, et al. (2020). Identification of a novel coronavirus causing severe pneumonia in human: a descriptive study. Chinese medical journal 133: 1015–1024.

4. Drosten, C, Gunther, S, Preiser, W, van der Werf, S, Brodt, HR, Becker, S, et al. (2003). Identification of a novel coronavirus in patients with severe acute respiratory syndrome. The New England journal of medicine 348: 1967–1976.

5. Zaki, AM, van Boheemen, S, Bestebroer, TM, Osterhaus, AD, and Fouchier, RA (2012). Isolation of a novel coronavirus from a man with pneumonia in Saudi Arabia. The New England journal of medicine 367: 1814–1820.

6. Zhu, N, Zhang, D, Wang, W, Li, X, Yang, B, Song, J, et al. (2020). A Novel Coronavirus from Patients with Pneumonia in China, 2019. The New England journal of medicine 382: 727–733.

7. Ksiazek, TG, Erdman, D, Goldsmith, CS, Zaki, SR, Peret, T, Emery, S, et al. (2003). A novel coronavirus associated with severe acute respiratory syndrome. The New England journal of medicine 348: 1953–1966.

8. Caly, L, Druce, JD, Catton, MG, Jans, DA, and Wagstaff, KM (2020). The FDA-approved drug ivermectin inhibits the replication of SARS-CoV-2 in vitro. Antiviral research 178: 104787.

9. Lobo-Galo, N, Terrazas-Lopez, M, Martinez-Martinez, A, and Diaz-Sanchez, AG (2020). FDA-approved thiol-reacting drugs that potentially bind into the SARS-CoV-2 main protease, essential for viral replication. Journal of biomolecular structure & dynamics: 1-9.

10. Jeon, S, Ko, M, Lee, J, Choi, I, Byun, SY, Park, S, et al. (2020). Identification of antiviral drug candidates against SARS-CoV-2 from FDA-approved drugs. Antimicrobial agents and chemotherapy.

11. Kandeel, M, and Al-Nazawi, M (2020). Virtual screening and repurposing of FDA approved drugs against COVID-19 main protease. Life sciences 251: 117627.

12. Ferner, RE, and Aronson, JK (2020). Chloroquine and hydroxychloroquine in covid-19. Bmj 369: m1432.

13. Singh, AK, Singh, A, Shaikh, A, Singh, R, and Misra, A (2020). Chloroquine and hydroxychloroquine in the treatment of COVID-19 with or without diabetes: A systematic search and a narrative review with a special reference to India and other developing countries. Diabetes & metabolic syndrome 14: 241–246.

14. Badyal, DK, and Mahajan, R (2020). Chloroquine: Can it be a Novel Drug for COVID-19. International journal of applied & basic medical research 10: 128–130.

15. Grein, J, Ohmagari, N, Shin, D, Diaz, G, Asperges, E, Castagna, A, et al. (2020). Compassionate Use of Remdesivir for Patients with Severe Covid-19. The New England journal of medicine.

16. Guastalegname, M, and Vallone, A (2020). Could chloroquine /hydroxychloroquine be harmful in Coronavirus Disease 2019 (COVID-19) treatment? Clinical infectious diseases : an official publication of the Infectious Diseases Society of America.

17. Gautret, P, Lagier, JC, Parola, P, Hoang, VT, Meddeb, L, Mailhe, M, et al. (2020). Hydroxychloroquine and azithromycin as a treatment of COVID-19: results of an open-label non-randomized clinical trial. International journal of antimicrobial agents: 105949.

18. Ou, X, Liu, Y, Lei, X, Li, P, Mi, D, Ren, L, et al. (2020). Characterization of spike glycoprotein of SARS-CoV-2 on virus entry and its immune cross-reactivity with SARS-CoV. Nature communications 11: 1620.

19. McIntosh, K, Dees, JH, Becker, WB, Kapikian, AZ, and Chanock, RM (1967). Recovery in tracheal organ cultures of novel viruses from patients with respiratory disease. Proceedings of the National Academy of Sciences of the United States of America 57: 933–940.

20. Wang, Y, Zhang, D, Du, G, Du, R, Zhao, J, Jin, Y, et al. (2020). Remdesivir in adults with severe COVID-19: a randomised, double-blind, placebo-controlled, multicentre trial. Lancet 395: 1569–1578.

21. Vinetz, JM (2020). Lack of efficacy of hydroxychloroquine in covid-19. Bmj 369: m2018.

22. Tang, W, Cao, Z, Han, M, Wang, Z, Chen, J, Sun, W, et al. (2020). Hydroxychloroquine in patients with mainly mild to moderate coronavirus disease 2019: open label, randomised controlled trial. Bmj 369: m1849.

23. Wang, M, Cao, R, Zhang, L, Yang, X, Liu, J, Xu, M, et al. (2020). Remdesivir and chloroquine effectively inhibit the recently emerged novel coronavirus (2019-nCoV) in vitro. Cell research 30: 269–271.

24. Woo, PC, Lau, SK, Chu, CM, Chan, KH, Tsoi, HW, Huang, Y, et al. (2005). Characterization and complete genome sequence of a novel coronavirus, coronavirus HKU1, from patients with pneumonia. Journal of virology 79: 884–895.

25. Zhang, Y, Li, J, Xiao, Y, Zhang, J, Wang, Y, Chen, L, et al. (2015). Genotype shift in human coronavirus OC43 and emergence of a novel genotype by natural recombination. The Journal of infection 70: 641–650.

26. Bao, L, Deng, W, Huang, B, Gao, H, Liu, J, Ren, L, et al. (2020). The pathogenicity of SARS-CoV-2 in hACE2 transgenic mice. Nature.

